# Modeling missing parents in single-step test-day SNP-BLUP evaluation of dairy cattle

**DOI:** 10.64898/2025.12.02.691779

**Authors:** Dawid Słomian, Jeremie Vandenplas, Jan Ten Napel, Kacper Żukowski, Monika Skarwecka, Joanna Szyda

## Abstract

In many countries, single-step genomic models have replaced multiple step models for routine evaluation. These models use all available information on animals’ phenotypes, genotypes, and pedigrees, yet missing parental information in pedigrees remains a challenge that affects genomic breeding value (GEBV) predictions. Therefore, the choice of method for handling missing parents can affect the prediction of breeding values. Here, we compared three approaches to model missing parental information for three levels of missing pedigree data: P_Real – pedigree from routine evaluation, P_2010 – at least 20 percent of dams and 10 percent of sires born before 2019 were set to missing, and P_4020 – at least 40 percent of dams and 20 percent of sires born before 2019 were set to missing. Missing parents’ information was expressed through missing codes in the raw pedigree (RP) by defining genetic groups (GG) that represent missing parents grouped based on year of birth, sex, and country of origin, or by defining metafounders (MF), which represent missing parents grouped by average genetic relationships estimated from the genomic information of their descendants. The genomic breeding values for fat yield were estimated using the single-step test-day SNP-BLUP model implemented with MiXBLUP software. For the considered scenarios, the results were presented separately for sires and dams, as well as for genotyped and ungenotyped individuals. We observed differences in the prediction quality between genotyped and ungenotyped animals. While GEBV predictions for the former were generally stable across scenarios, the predictions for the ungenotyped individuals varied. In particular, the removal of parental information led to less stable results when missing parental information was expressed by MF, where insufficient pedigree completeness resulted in an overestimation of the genetic trend. In conclusion, for informative pedigrees with a small percentage of missing parents, the incorporation of GG and MF results in very similar GEBV predictions, however GG appear to be a more robust approach for ungenotyped individuals in highly incomplete pedigrees.

## Introduction

Currently, many countries implement a single-step model that includes all available animal information (phenotype, genotype, and pedigree) for routine genetic evaluation. The structure of pedigree data is an important aspect of the genetic evaluation of dairy cattle (Bradford et al., 2019). Therefore, a practical challenge is to deal with missing individuals in the pedigree. The standard approach for this situation is to combine missing individuals within unrelated genetic groups (or phantom parent groups), which are typically defined based on the country of origin, sex, and year of birth (Westell et al., 1988; Legarra et al., 2007). Although genetic groups are commonly used, they can lead to bias and excessive scattering of genomically estimated breeding values (GEBV), especially if there are many missing parents (Masuda et al., 2021; Himmelbauer et al., 2024). Defining metafounders (Legarra et al., 2015) is a genotype-aware alternative to genetic groups. This approach involves forming groups of missing parents based on average genetic relationships estimated from single nucleotide polymorphisms (SNPs) information of their descendants, for which a 0.5 allele frequency at all SNPs is assumed. However, Kudinov et al. (2020) did not observe any differences in GEBVs predicted using genetic groups and metafounder approaches to combine missing parents. On the other hand Bradford et al. (2019), Macedo et al. (2020), Kudinov et al. (2022), and Himmelbauer et al. (2024) reported more accurate predictions of breeding values when using metafounders as compared to genetic groups, especially in pedigrees with low pedigree completeness and for low heritable traits.

As all the above-mentioned studies compared the use of genetic groups and metafounders for GEBV prediction based on the G-BLUP model, we aimed to compare the approaches of handling missing pedigree in the single-step SNP-BLUP formulation. In particular, three levels of missing parent data and three approaches for handling them were considered. The comparison was applied to a real dataset representing the national evaluation for fat yield. In particular, we focused on the GEBV validation results, GEBV trends, and accuracy of GEBV prediction for young individuals.

## Material

The data originated from the Polish national evaluation run for fat yield from April 2024 (Table 1) and included 3,707,727 cows with 63,615,019 records in the full dataset and 3,224,917 cows with 58,446,695 records in the truncated dataset, defined by removing animals born after 2018. Genotypic information was available for 181,991 individuals and comprised 46,118 SNP genotypes. The pedigree used in this study contained 4,712,143 animals and originated from the raw pedigree file by extracting individuals representing the third generation before the oldest individual with data (phenotype or genotype) using the RelaX2 software (Strandén, 2014).

**Table 1.** Number of animals in the analyzed dataset.

| Data | Sex | Number of<br>animals | Number of<br>records |
| --- | --- | --- | --- |
| Phenotype Full dataset (fat yield) | Cows | 3,707,727 | 63,615,019 |
| Phenotype Truncated dataset (fat yield) |  | 3,224,917 | 58,446,695 |
| Genotype | Cows | 113,019 | 181,991 |
|  | Bulls | 68,972 |  |
| Pedigree | Cows | 4,569,044 | 4,712,143 |
|  | Bulls | 143,099 |  |

Based on this pedigree, the following scenarios were considered:

- Pedigree_real (**P_Real**) – the original pedigree from routine evaluation containing 262,519 (5.6%) missing sires and 719,360 (15.3%) missing dams,
- Pedigree_20_10 (**P_2010**) – **P_real** with approximately 20% of the dams and 10% of the sires set to missing, containing 446,669 (9.5%) missing bulls and 1,076,127 (22.8%) missing cows,
- Pedigree_40_20 **(P_4020**) – **P_real** with approximately 40% of the dams and 20% of the sires set to missing, containing 884,192 (18.7%) missing bulls and 1,868,957 (39.6%) missing cows.

Note that the pedigree of the youngest animals, born from 2019, that were used for validation was the same in all scenarios. Differences between scenarios, therefore, start with the pedigree of the second generation, that is, the parents of these young individuals (Figure 1). Moreover, only parental information was removed, so that the phenotypic records of the parents remained in the data.

**Figure 1.**
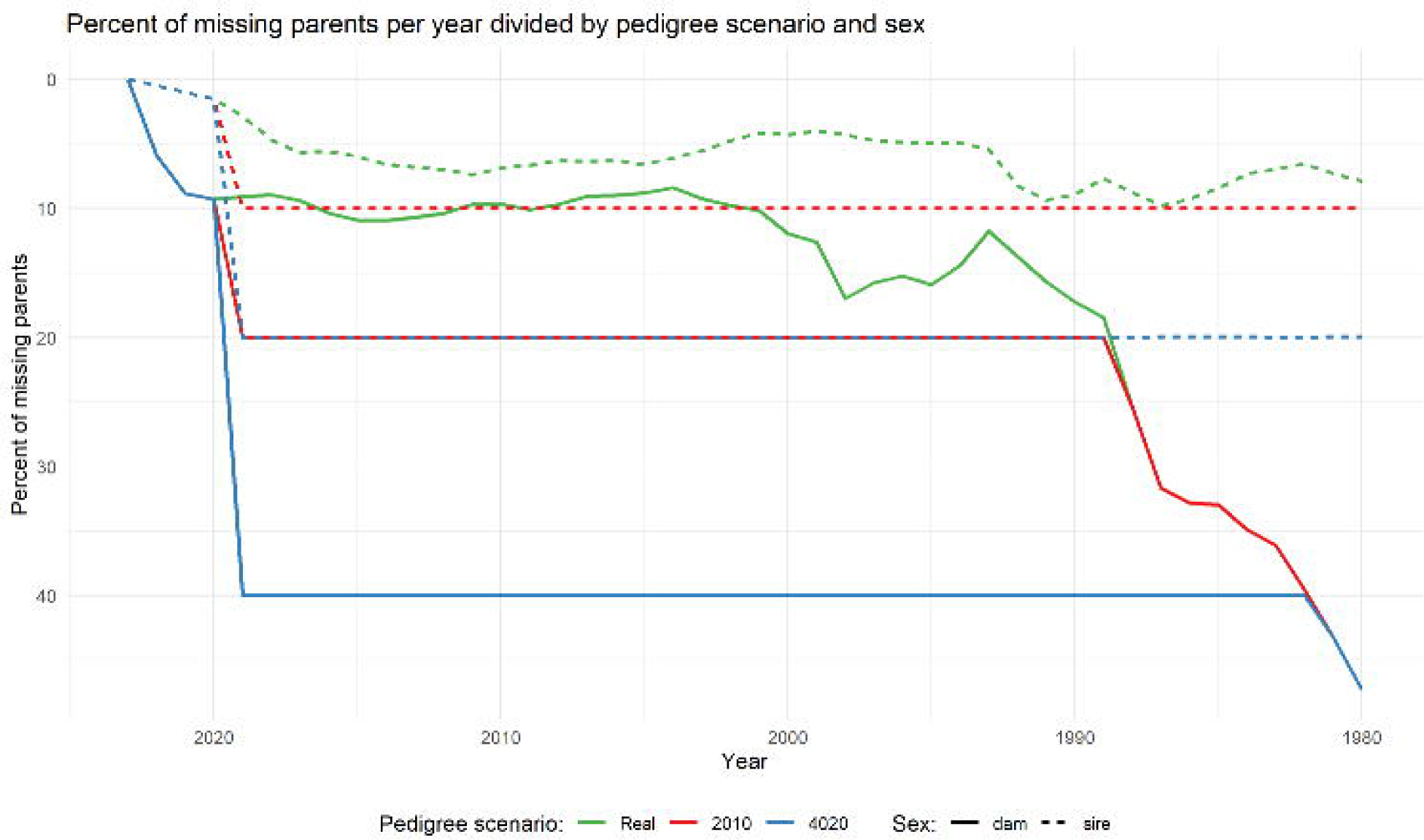
Percentage of missing parents for each pedigree scenario.

Furthermore, for the defined pedigrees, three approaches for handling missing parental information were implemented:

- Raw pedigree (**RP**) – missing parents’ IDs set to missing,
- Genetic groups (**GG**) – missing parents replaced by unrelated **GG** defined based on birth year (10-year grouping starting from 1960, with individuals born before 1960 assigned to a common genetic group), country of origin (Poland, USA, Canada, other), and sex,
- Metafounders (**MF**) – missing parents replaced by **MF**, which represent genetic groups with relationships estimated from the genomic information of descendants.

The number of defined **GGs** differed between pedigree scenarios. For **P_2010** and **P_4020,** more **GGs** were defined than for **P_Real**. The highest rate of missing parents in each scenario was observed for animals born in Poland between 1990 and 2019 (Figure 2). Since **MF** were created based on **GG**, the numbers of **MF** and **GG** were the same.

**Figure 2.**
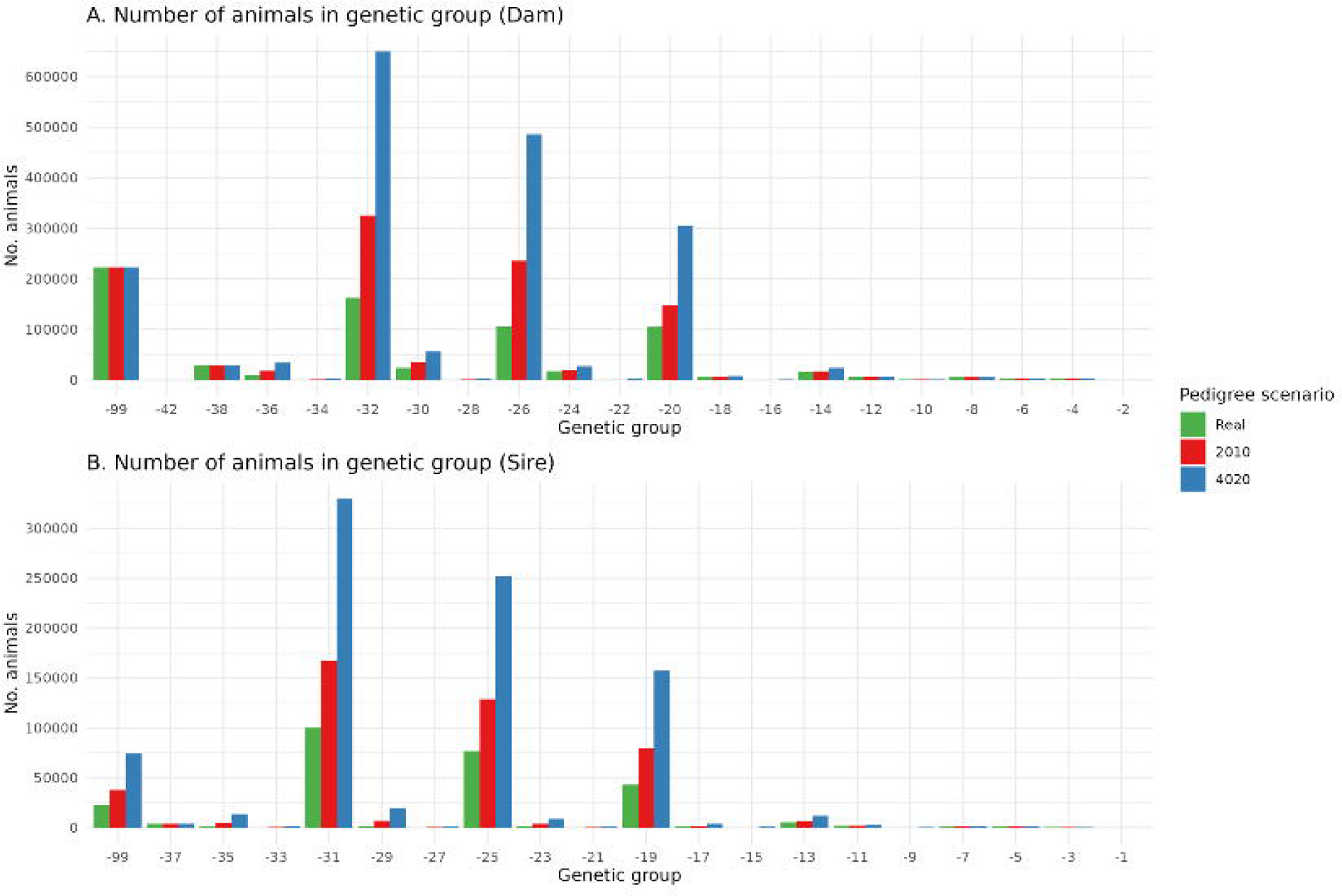
Number of individuals in each genetic group for each pedigree scenario.

## Methods

The following single-step test-day SNP-BLUP model (Liu et al., 2004) was used to predict breeding values:

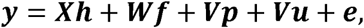

where ***y*** contains cow test day records for fat yield from the first three parities, ***h*** is a vector of fixed effects of herd-test_date-parity-milking_frequency, ***f*** is a vector of fixed lactation curve coefficients, which was modeled by the Wilmink function (Liu et al., 2004), ***p*** is a vector of permanent environmental effects expressed as three random regression coefficients of the Legendre polynomial, and ***u*** is a random additive genetic effect also described by the three random regression coefficients of the Legendre polynomials. ***V*** is a design matrix for ***u*, *p*** contains the Legendre coefficients for the first three lactations, ***X*** is the design matrix for the fixed herd-test-date-parity-milking-frequency effect ***h***, and ***W*** is the design matrix for fixed effect ***f***, and 𝒆 is a residual. The model was implemented using MiXBLUP 3.0 (Vandenplas et al., 2022).

The validation of 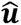 was conducted on 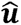 representing the 305-day Genomically Enhanced Breeding Values (GEBVs) combined over all three lactations, using the following pattern 𝑮𝑬𝑩𝑽𝑡 = 0.5𝑮𝑬𝑩𝑽1 + 0.3𝑮𝑬𝑩𝑽2 + 0.2𝑮𝑬𝑩𝑽3, where ***GEBV****t* is the combined GEBV and ***GEBV****i* represents GEBV for the i-th lactation. The validation test conducted following Mäntysaari et al. (2010) was implemented separately for cows and bulls based on the standardized ***GEBV****t* as:

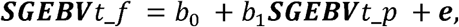

where ***SGEBV****t_f* represents the vector of standardized GEBVs predicted based on the full dataset, while ***SGEBV****t_p* represents standardized GEBVs predicted based on the truncated dataset, *b_0_* represents the intercept, which indicates a systematic bias in the model’s prediction, and *b_1_* represents the regression slope, that is, the divergence between predictions and actual GEBVs. The R^2^ coefficient that quantifies the percentage of variance of ***SGEBV****t_f* explained by 𝑺𝑮𝑬𝑩𝑽𝑡_𝑝 was used as a measure of prediction accuracy. Validation cows were defined as dams born between 2019 and 2022 with test-day records for at least one lactation. Validation bulls were defined as sires born between 2017 and 2020 with at least 20 daughters. The standardization of GEBVs was performed separately for the full and truncated datasets as: 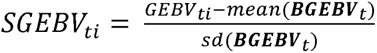, where 𝑩𝑮𝑬𝑩𝑽_𝑡_ denotes the vector of GEBVs of base animals represented by cows with phenotypes.

Figure 3 shows the average differences in the number of progeny of the validation bulls between **P_2010** and **P_4020** scenarios relative to **P_Real**. The average difference in the number of progeny for **P_2010** was 4 (sires born in 2017), 2 (sires born in 2018), and 7 (sires born in 2019) for genotyped daughters and 1 (sires born in 2017), 2 (sires born in 2018), and 2 (sires born in 2019) for ungenotyped daughters, respectively. For **P_4020,** the differences were 10 (sires born in 2017), 5 (sires born in 2018), and 22 (sires born in 2019) for genotyped daughters and 3 (sires born in 2017), 4 (sires born in 2018), and 4 (sires born in 2019) for ungenotyped daughters. The highest mean difference in the number of sons was observed for 2017 for **P_4020**, genotyped bulls (1 son), whereas for ungenotyped bulls, for **P_2010** and **P_4020**, it was 0 for 2019.

**Figure 3.**
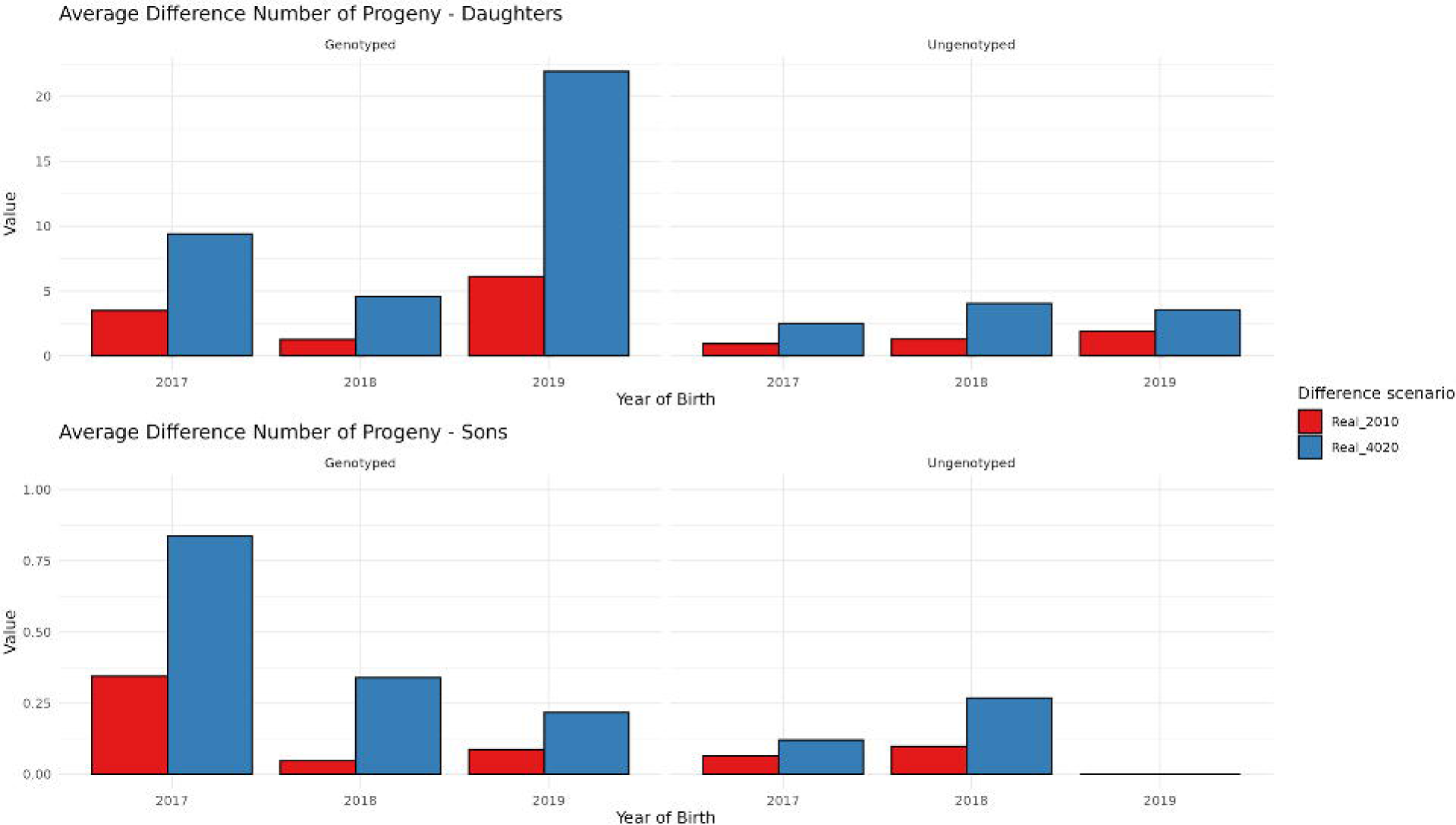
Average difference in the number of progenies between RP and 2010, 4020 divided by genotyped and ungenotyped individuals.

## Results

### Validation

The validation model was computed for 562 validation bulls (387 genotyped and 175 ungenotyped) and 482,810 validation cows (30,227 genotyped and 452,336 ungenotyped). Figure 4A shows the estimated slopes (b1) of the validation models divided by scenario, sex, parity, and genotyping status. For bulls, the value of b_1_ was close to the expected value of one for most scenarios, except **P_2010** (1.271) and **P_4020** (1.328) for **MF** for ungenotyped bulls. Moreover, for MF, overdispersion was observed when comparing the predictions of P_2010 and P_4020 with P_Real. Figure 4B shows intercepts (b_0_) for all scenarios. Similar estimates, close to zero, were observed for every scenario, where the minimum value was −0.463 (**P_2010** for **RP** for genotyped bulls) and the maximum value was 0.067 (**P_4020** for **MF** for genotyped cows). In general, the intercepts estimated for bulls were negative, whereas for cows, the values were very close to zero. Figure 4C shows R^2^,and Figure 4D displays the Pearson correlation coefficients between GEBVs predicted from the truncated and full datasets for the validation animals, divided by scenario, sex, parity, and genotyping status. When **MF** or **GG** were used, the R² and correlations for cows were generally higher than those for the bulls. Additionally, the GEBV of genotyped cows always resulted in higher correlations than those of ungenotyped cows, regardless of missing parent handling. However, for **P_Real**, R^2^ and correlations depended on the adopted missing parent approach. The most striking scenario was for the ungenotyped **P_Real**, where the correlation for **RP** was 0.892, decreased to 0.870 for **GG**, but increased to 0.924 for **MF**. The correlations for bulls followed a different pattern. For ungenotyped bulls, the correlation in each case increased sequentially from **RP** through **GG** to **MF**. For genotyped bulls, for **P_Real,** the correlation was similar across scenarios, but for **P_2010** and **P_4020, MF** resulted in slightly lower correlations.

**Figure 4.**
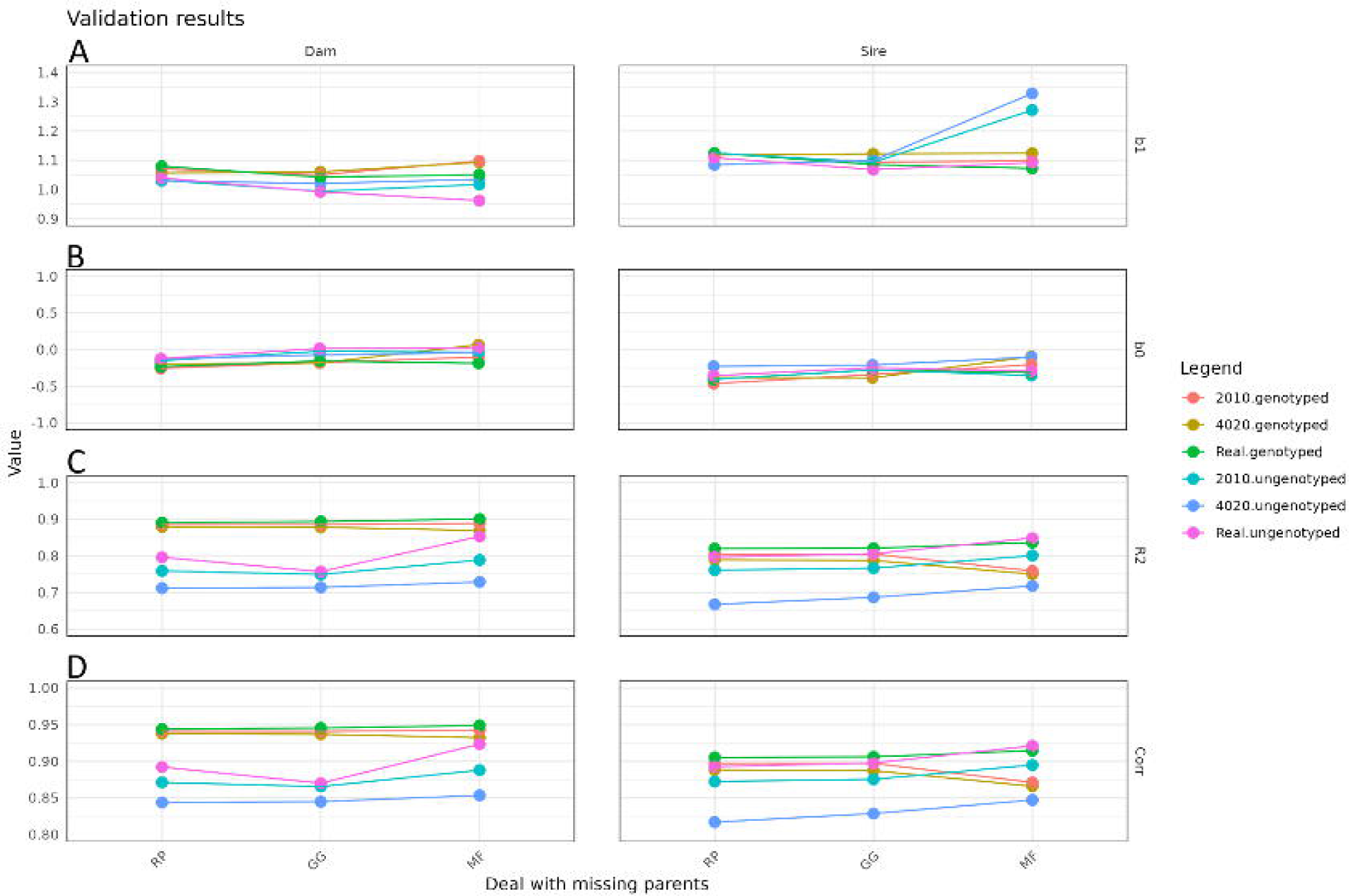
Validation results for individuals divided by sex and method.

Tables A1-A4 in the appendix provide detailed validation results for the genotyped and ungenotyped validation bulls and cows, respectively.

### GEBV comparison

Figures 5 and 6 show GEBVs predicted for the full and the truncated datasets for dams and sires. For each scenario, the points on the scatter plots were grouped into elliptical clusters along the diagonal. However, for ungenotyped individuals, especially dams, we observed greater dispersion along the diagonal line. For ungenotyped bulls under the **P_4020** scenarios, GEBVs predicted from the truncated dataset were underestimated for bulls with the highest GEBSs in the full data, regardless of the applied approach of missing parent handling. Figure 7 shows the average GEBV trends for all scenarios divided by sex. The average GEBV trend is a combination of the population average (genetic trend: year-to-year change in the mean breeding value) and the intensity of selection (the advantage of selected parents within the year). After 2010, we observed that the mean GEBV increased for all animals, especially bulls. For each scenario, the highest increase in the GEBV trend was realized by **P_Real** with missing parents handled via **MF.**

**Figure 5.**
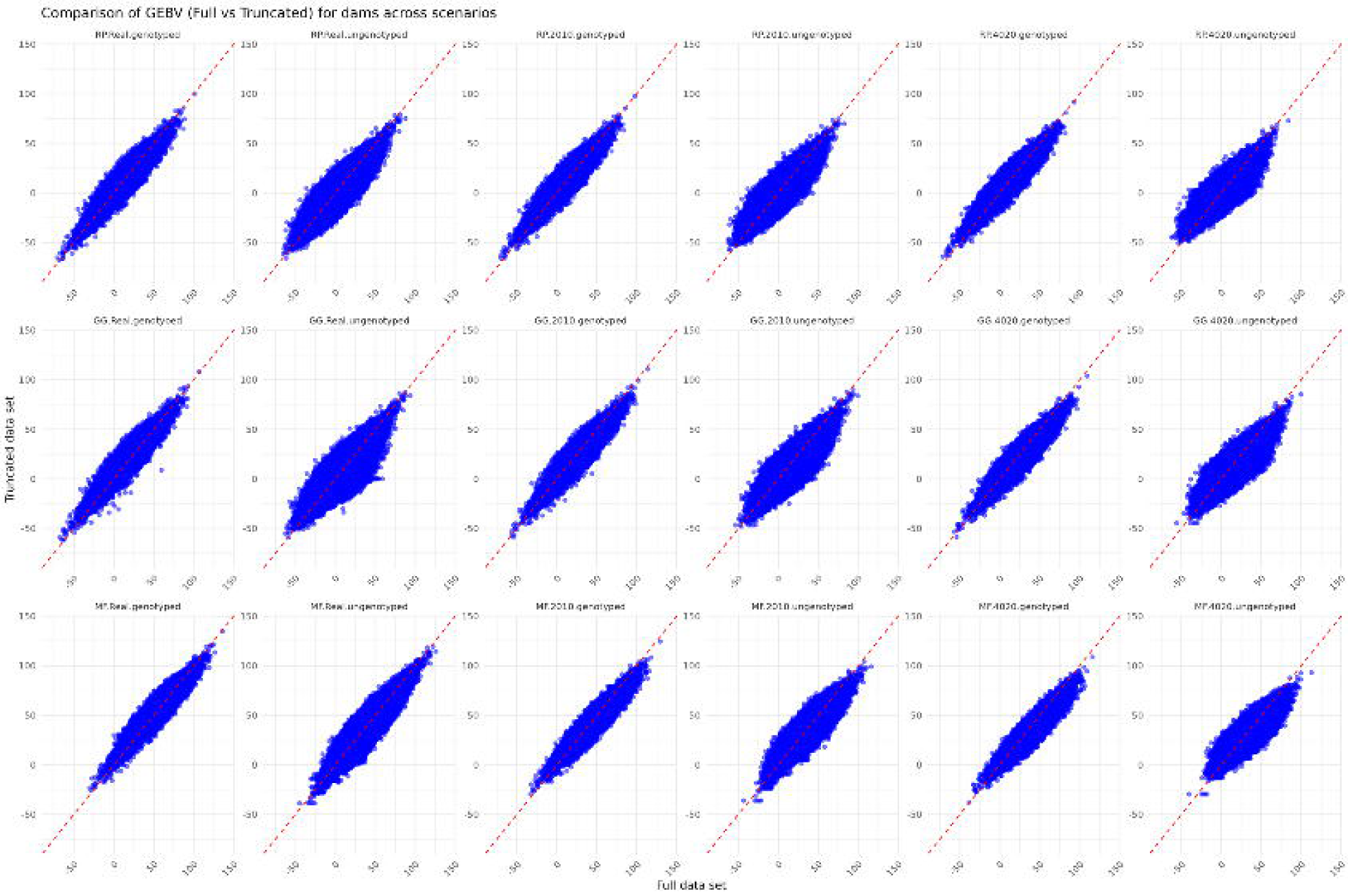
Comparison of GEBV for dams across scenarios divided into genotyped and ungenotyped individuals.

**Figure 6.**
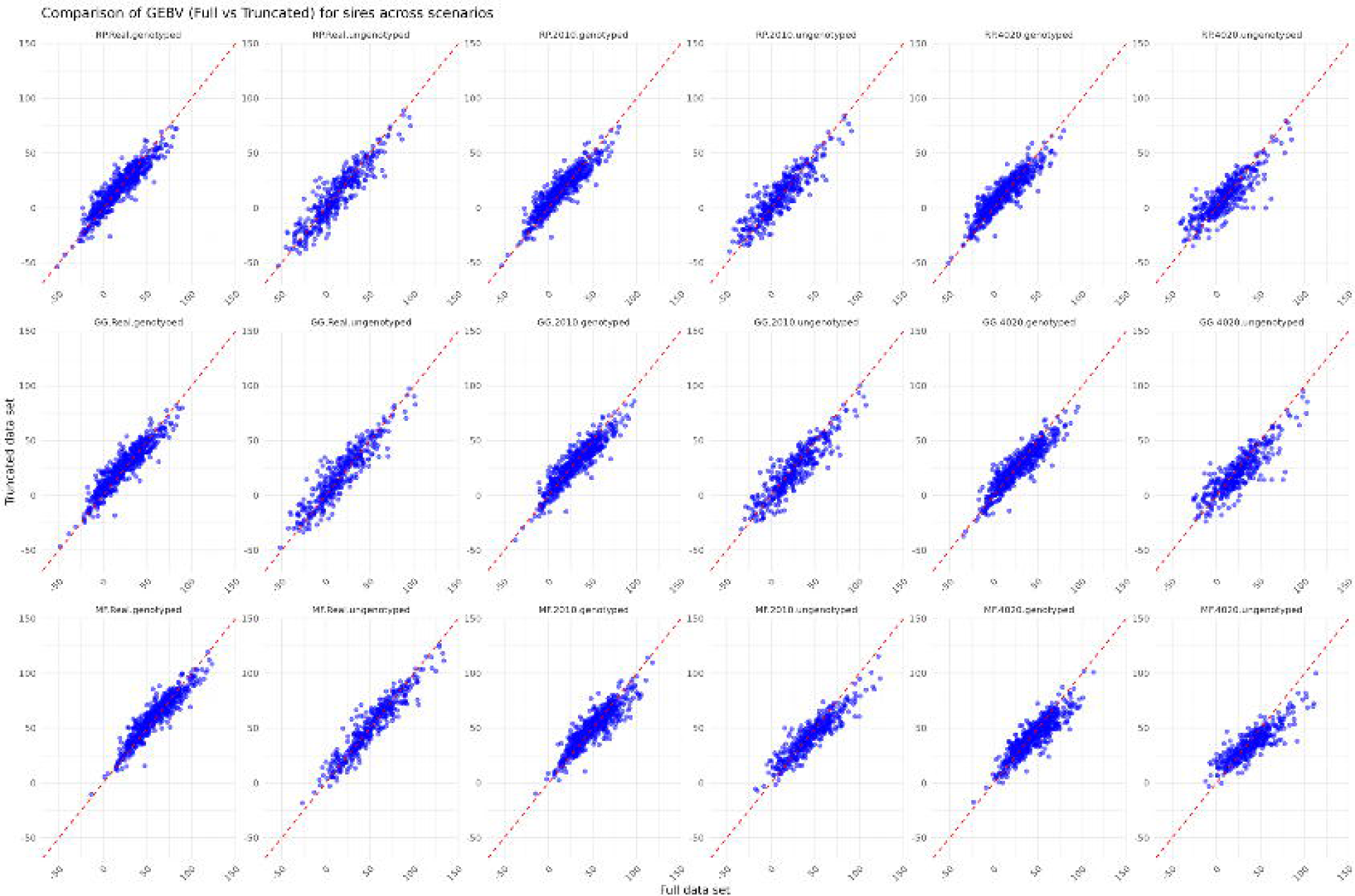
Comparison of GEBV for sires across scenarios divided into genotyped and ungenotyped individuals.

**Figure 7.**
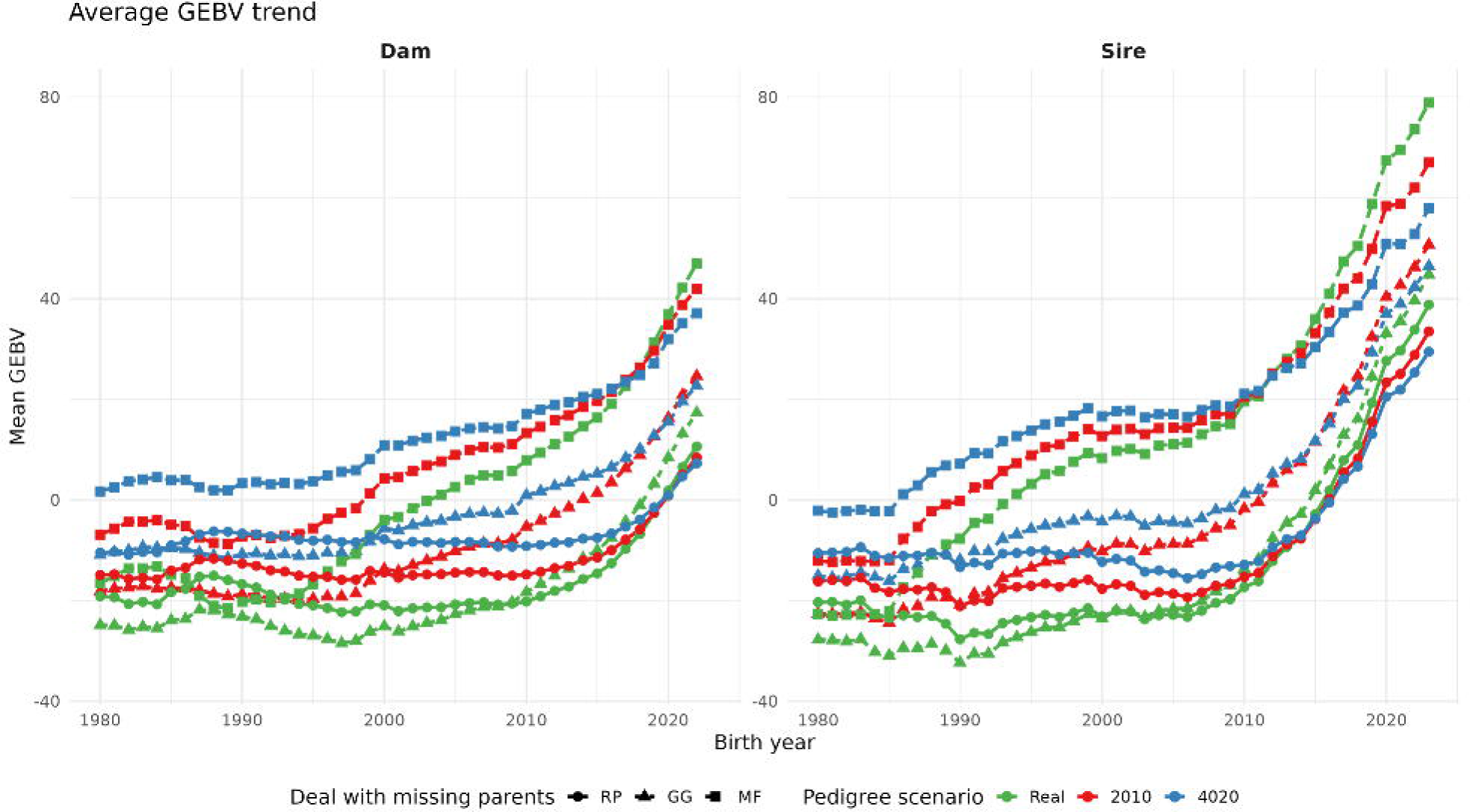
Genetic trends for individuals divided by sex and method.

## Discussion

Our study focused on the comparison of different methods for expressing missing parental information (**RP**, **GG**, or **MF**) in the context of a single-step evaluation using the SNP-BLUP test-day model. In addition, we investigated the robustness of these three approaches to the increasing amount of missing parental information in the pedigree, separately for the genotyped and ungenotyped parts of the population. The analysis was conducted based on the real pedigree representing the Polish dairy cattle population, using fat yield as an example, and was assessed in terms of the statistical properties underlying the GEBV validation model. Two aspects emerge from these comparisons. First, the availability of genotype information enhances the quality of GEBVs prediction not only by resulting in “better” estimates of the parameters of the regression line in validation but also in higher R^2^ and predicted versus observed GEBVs’ correlations, as compared to ungenotyped individuals. Owing to the availability of genomic information, the choice of the method of missing parent handling does not impact GEBV prediction quality. The second aspect demonstrated in our study is that for ungenotyped individuals, the **MF** approach is not robust with respect to the number of missing parents. As the percentage of metafounders increases, it struggles to pass genotypic information to ungenotyped individuals. In particular, we observed that using metafounders led to the systematic underestimation of the GEBVs of the ungenotyped bulls with an increasing number of incomplete pedigree data, and consequently to an elevated b_1_ in the validation model. Some, albeit not very strong, advantages of using **MF** over **GG** were previously reported, in the context of a single step G-BLUP evaluation, by Garcia-Baccino et al. (2017), Macedo et al. (2020), Macedo et al. (2022), and Kluska et al. (2021). The first study was based on simulated data and reported good predictive performance using **MF**. For the dairy sheep population, Macedo et al. (2020, 2022) obtained the most accurate GEBV prediction results, expressed by a validation slope close to unity, with **MF**. Kluska et al. (2021) reported that in a beef cattle population, using **MF** and **GG** provided very similar results, but finally **MF** was ultimately recommended as the approach providing the smallest bias of GEBVs.

## Conclusions

In summary, this study demonstrates that methods for handling missing parents in pedigrees may impact GEBV and that handling missing parents is increasingly important with the increasing number of incomplete pedigrees. The most important result of this study is that using the metafounder approach may lead to biased predictions for ungenotyped individuals, particularly as the proportion of missing parents increases. In contrast, for genotyped individuals, no marked differences in the handling of missing parent data were observed.

