## Appendix A for "Modeling missing parents in single-step test-day SNP-BLUP evaluation of dairy cattle"

Table A1 Validation results for genotyped validation bulls (n=387).

| Value | Deal with missing parents | Pedigree Scenario | 1^st^ lactation | 2^nd^ lactation | 3^rd^ lactation | Total GEBV |
| --- | --- | --- | --- | --- | --- | --- |
| b_0_ | RP | Real | -0.43 | -0.39 | -0.29 | -0.40 |
|  |  | 2010 | -0.47 | -0.48 | -0.37 | -0.46 |
|  |  | 4020 | -0.39 | -0.38 | -0.31 | -0.38 |
|  | GG | Real | -0.28 | -0.29 | -0.22 | -0.28 |
|  |  | 2010 | -0.33 | -0.37 | -0.27 | -0.34 |
|  |  | 4020 | -0.37 | -0.41 | -0.32 | -0.38 |
|  | MF | Real | -0.30 | -0.34 | -0.27 | -0.31 |
|  |  | 2010 | -0.19 | -0.22 | -0.18 | -0.20 |
|  |  | 4020 | -0.08 | -0.09 | -0.07 | -0.09 |
| b_1_ | RP | Real | 1.13 | 1.13 | 1.10 | 1.12 |
|  |  | 2010 | 1.11 | 1.14 | 1.10 | 1.12 |
|  |  | 4020 | 1.10 | 1.13 | 1.11 | 1.12 |
|  | GG | Real | 1.08 | 1.09 | 1.07 | 1.08 |
|  |  | 2010 | 1.09 | 1.10 | 1.07 | 1.09 |
|  |  | 4020 | 1.12 | 1.13 | 1.10 | 1.12 |
|  | MF | Real | 1.07 | 1.08 | 1.06 | 1.07 |
|  |  | 2010 | 1.10 | 1.10 | 1.08 | 1.10 |
|  |  | 4020 | 1.13 | 1.12 | 1.10 | 1.12 |
| R^2^ | RP | Real | 0.80 | 0.83 | 0.85 | 0.82 |
|  |  | 2010 | 0.79 | 0.81 | 0.83 | 0.80 |
|  |  | 4020 | 0.78 | 0.79 | 0.81 | 0.79 |
|  | GG | Real | 0.81 | 0.83 | 0.85 | 0.82 |
|  |  | 2010 | 0.79 | 0.81 | 0.83 | 0.80 |
|  |  | 4020 | 0.77 | 0.79 | 0.82 | 0.79 |
|  | MF | Real | 0.82 | 0.85 | 0.86 | 0.84 |
|  |  | 2010 | 0.75 | 0.76 | 0.79 | 0.76 |
|  |  | 4020 | 0.74 | 0.76 | 0.78 | 0.75 |
| Corr. | RP | Real | 0.90 | 0.91 | 0.92 | 0.91 |
|  |  | 2010 | 0.89 | 0.90 | 0.91 | 0.90 |
|  |  | 4020 | 0.88 | 0.89 | 0.90 | 0.89 |
|  | GG | Real | 0.90 | 0.91 | 0.92 | 0.91 |
|  |  | 2010 | 0.89 | 0.90 | 0.91 | 0.90 |
|  |  | 4020 | 0.88 | 0.89 | 0.90 | 0.89 |
|  | MF | Real | 0.91 | 0.92 | 0.93 | 0.91 |
|  |  | 2010 | 0.86 | 0.87 | 0.89 | 0.87 |
|  |  | 4020 | 0.86 | 0.87 | 0.88 | 0.87 |

b_0_ – slope
b_1_ – intercept
R^2^ – coefficient of determination
Corr. - Correlation

Table A2 Validation results for ungenotyped validation bulls (n=175).

| Value | Deal with missing parents | Pedigree Scenario | 1^st^ lactation | 2^nd^ lactation | 3^rd^ lactation | Total GEBV |
| --- | --- | --- | --- | --- | --- | --- |
| b_0_ | RP | Real | -0.37 | -0.38 | -0.25 | -0.35 |
|  |  | 2010 | -0.40 | -0.46 | -0.30 | -0.41 |
|  |  | 4020 | -0.20 | -0.30 | -0.15 | -0.23 |
|  | GG | Real | -0.25 | -0.28 | -0.18 | -0.25 |
|  |  | 2010 | -0.26 | -0.32 | -0.21 | -0.28 |
|  |  | 4020 | -0.16 | -0.30 | -0.15 | -0.21 |
|  | MF | Real | -0.29 | -0.32 | -0.24 | -0.29 |
|  |  | 2010 | -0.33 | -0.38 | -0.28 | -0.35 |
|  |  | 4020 | -0.04 | -0.15 | -0.06 | -0.10 |
| b_1_ | RP | Real | 1.12 | 1.10 | 1.08 | 1.11 |
|  |  | 2010 | 1.13 | 1.13 | 1.09 | 1.12 |
|  |  | 4020 | 1.08 | 1.10 | 1.05 | 1.08 |
|  | GG | Real | 1.08 | 1.07 | 1.05 | 1.07 |
|  |  | 2010 | 1.10 | 1.09 | 1.06 | 1.09 |
|  |  | 4020 | 1.11 | 1.10 | 1.05 | 1.10 |
|  | MF | Real | 1.10 | 1.09 | 1.07 | 1.09 |
|  |  | 2010 | 1.29 | 1.26 | 1.21 | 1.27 |
|  |  | 4020 | 1.34 | 1.32 | 1.26 | 1.33 |
| R^2^ | RP | Real | 0.78 | 0.81 | 0.82 | 0.80 |
|  |  | 2010 | 0.74 | 0.78 | 0.78 | 0.76 |
|  |  | 4020 | 0.64 | 0.70 | 0.69 | 0.67 |
|  | GG | Real | 0.79 | 0.82 | 0.82 | 0.81 |
|  |  | 2010 | 0.75 | 0.78 | 0.79 | 0.77 |
|  |  | 4020 | 0.66 | 0.72 | 0.71 | 0.69 |
|  | MF | Real | 0.83 | 0.86 | 0.86 | 0.85 |
|  |  | 2010 | 0.78 | 0.81 | 0.81 | 0.80 |
|  |  | 4020 | 0.70 | 0.74 | 0.73 | 0.72 |
| Corr. | RP | Real | 0.88 | 0.90 | 0.91 | 0.89 |
|  |  | 2010 | 0.86 | 0.88 | 0.89 | 0.87 |
|  |  | 4020 | 0.80 | 0.84 | 0.83 | 0.82 |
|  | GG | Real | 0.89 | 0.90 | 0.91 | 0.90 |
|  |  | 2010 | 0.86 | 0.88 | 0.89 | 0.88 |
|  |  | 4020 | 0.81 | 0.85 | 0.84 | 0.83 |
|  | MF | Real | 0.91 | 0.93 | 0.93 | 0.92 |
|  |  | 2010 | 0.89 | 0.90 | 0.90 | 0.89 |
|  |  | 4020 | 0.84 | 0.86 | 0.85 | 0.85 |

b_0_ – slope
b_1_ – intercept
R^2^ – coefficient of determination
Corr. - Correlation

Table A3 Validation results for genotyped validation cows (n=30,227).

| Value | Deal with missing parents | Pedigree Scenario | 1^st^ lactation | 2^nd^ lactation | 3^rd^ lactation | Total GEBV |
| --- | --- | --- | --- | --- | --- | --- |
| b_0_ | RP | Real | -0.25 | -0.24 | -0.16 | -0.23 |
|  |  | 2010 | -0.26 | -0.28 | -0.20 | -0.26 |
|  |  | 4020 | -0.19 | -0.21 | -0.16 | -0.19 |
|  | GG | Real | -0.15 | -0.17 | -0.12 | -0.15 |
|  |  | 2010 | -0.17 | -0.21 | -0.15 | -0.18 |
|  |  | 4020 | -0.16 | -0.22 | -0.16 | -0.18 |
|  | MF | Real | -0.18 | -0.20 | -0.16 | -0.19 |
|  |  | 2010 | -0.09 | -0.11 | -0.09 | -0.10 |
|  |  | 4020 | 0.08 | 0.06 | 0.06 | 0.07 |
| b_1_ | RP | Real | 1.09 | 1.08 | 1.06 | 1.08 |
|  |  | 2010 | 1.07 | 1.08 | 1.05 | 1.07 |
|  |  | 4020 | 1.05 | 1.07 | 1.05 | 1.06 |
|  | GG | Real | 1.04 | 1.05 | 1.03 | 1.04 |
|  |  | 2010 | 1.05 | 1.06 | 1.03 | 1.05 |
|  |  | 4020 | 1.06 | 1.07 | 1.05 | 1.06 |
|  | MF | Real | 1.05 | 1.06 | 1.04 | 1.05 |
|  |  | 2010 | 1.10 | 1.10 | 1.08 | 1.10 |
|  |  | 4020 | 1.10 | 1.09 | 1.08 | 1.09 |
| R^2^ | RP | Real | 0.88 | 0.90 | 0.91 | 0.89 |
|  |  | 2010 | 0.87 | 0.89 | 0.90 | 0.89 |
|  |  | 4020 | 0.87 | 0.88 | 0.89 | 0.88 |
|  | GG | Real | 0.88 | 0.90 | 0.91 | 0.89 |
|  |  | 2010 | 0.88 | 0.89 | 0.90 | 0.89 |
|  |  | 4020 | 0.87 | 0.88 | 0.89 | 0.88 |
|  | MF | Real | 0.89 | 0.91 | 0.91 | 0.90 |
|  |  | 2010 | 0.88 | 0.89 | 0.90 | 0.89 |
|  |  | 4020 | 0.86 | 0.87 | 0.88 | 0.87 |
| Corr. | RP | Real | 0.94 | 0.95 | 0.95 | 0.94 |
|  |  | 2010 | 0.94 | 0.94 | 0.95 | 0.94 |
|  |  | 4020 | 0.93 | 0.94 | 0.95 | 0.94 |
|  | GG | Real | 0.94 | 0.95 | 0.95 | 0.95 |
|  |  | 2010 | 0.94 | 0.94 | 0.95 | 0.94 |
|  |  | 4020 | 0.93 | 0.94 | 0.95 | 0.94 |
|  | MF | Real | 0.94 | 0.95 | 0.95 | 0.95 |
|  |  | 2010 | 0.94 | 0.94 | 0.95 | 0.94 |
|  |  | 4020 | 0.93 | 0.93 | 0.94 | 0.93 |

b_0_ – slope
b_1_ – intercept
R^2^ – coefficient of determination
Corr. - Correlation

Table A4 Validation results for ungenotyped validation cows (n=452,336).

| Value | Deal with missing parents | Pedigree Scenario | 1^st^ lactation | 2^nd^ lactation | 3^rd^ lactation | Total GEBV |
| --- | --- | --- | --- | --- | --- | --- |
| b_0_ | RP | Real | -0.14 | -0.13 | -0.09 | -0.12 |
|  |  | 2010 | -0.15 | -0.17 | -0.12 | -0.15 |
|  |  | 4020 | -0.12 | -0.15 | -0.11 | -0.13 |
|  | GG | Real | 0.03 | -0.01 | 0.02 | 0.02 |
|  |  | 2010 | -0.01 | -0.05 | -0.02 | -0.02 |
|  |  | 4020 | -0.05 | -0.11 | -0.07 | -0.07 |
|  | MF | Real | 0.04 | 0.01 | 0.02 | 0.03 |
|  |  | 2010 | -0.02 | -0.06 | -0.04 | -0.04 |
|  |  | 4020 | -0.02 | -0.06 | -0.04 | -0.04 |
| b_1_ | RP | Real | 1.04 | 1.04 | 1.04 | 1.04 |
|  |  | 2010 | 1.03 | 1.04 | 1.03 | 1.03 |
|  |  | 4020 | 1.02 | 1.04 | 1.03 | 1.03 |
|  | GG | Real | 0.98 | 1.00 | 1.00 | 0.99 |
|  |  | 2010 | 0.99 | 1.00 | 1.00 | 0.99 |
|  |  | 4020 | 1.01 | 1.03 | 1.02 | 1.02 |
|  | MF | Real | 0.96 | 0.97 | 0.97 | 0.96 |
|  |  | 2010 | 1.01 | 1.02 | 1.02 | 1.02 |
|  |  | 4020 | 1.03 | 1.04 | 1.04 | 1.03 |
| R^2^ | RP | Real | 0.78 | 0.80 | 0.82 | 0.80 |
|  |  | 2010 | 0.75 | 0.77 | 0.78 | 0.76 |
|  |  | 4020 | 0.70 | 0.72 | 0.74 | 0.71 |
|  | GG | Real | 0.73 | 0.77 | 0.79 | 0.76 |
|  |  | 2010 | 0.73 | 0.76 | 0.78 | 0.75 |
|  |  | 4020 | 0.70 | 0.73 | 0.74 | 0.71 |
|  | MF | Real | 0.84 | 0.86 | 0.87 | 0.85 |
|  |  | 2010 | 0.78 | 0.80 | 0.81 | 0.79 |
|  |  | 4020 | 0.72 | 0.74 | 0.75 | 0.73 |
| Corr. | RP | Real | 0.88 | 0.90 | 0.90 | 0.89 |
|  |  | 2010 | 0.86 | 0.88 | 0.88 | 0.87 |
|  |  | 4020 | 0.84 | 0.85 | 0.86 | 0.84 |
|  | GG | Real | 0.86 | 0.88 | 0.89 | 0.87 |
|  |  | 2010 | 0.85 | 0.87 | 0.88 | 0.87 |
|  |  | 4020 | 0.83 | 0.85 | 0.86 | 0.84 |
|  | MF | Real | 0.92 | 0.93 | 0.93 | 0.92 |
|  |  | 2010 | 0.88 | 0.89 | 0.90 | 0.89 |
|  |  | 4020 | 0.85 | 0.86 | 0.87 | 0.85 |

b_0_ – slope
b_1_ – intercept
R^2^ – coefficient of determination
Corr. - Correlation
